# Functional brain age prediction suggests accelerated aging in preclinical familial Alzheimer’s disease, irrespective of fibrillar amyloid-beta pathology

**DOI:** 10.1101/2020.05.06.076745

**Authors:** Julie Gonneaud, Alex T. Baria, Alexa Pichet Binette, Brian A. Gordon, Jasmeer P. Chhatwal, Carlos Cruchaga, Mathias Jucker, Johannes Levin, Stephen Salloway, Martin Farlow, Serge Gauthier, Tammie L.S. Benzinger T, John C. Morris, Randall J. Bateman, John C.S. Breitner, Judes Poirier, Etienne Vachon-Presseau, Sylvia Villeneuve, for the Alzheimer’s Disease Neuroimaging Initiative, the Dominantly Inherited Alzheimer Network (DIAN), the PREVENT-AD Research Group

## Abstract

We aimed at developing a model able to predict brain aging from resting state functional connectivity (rs-fMRI) and assessing whether genetic risk/determinants of Alzheimer’s disease (AD) and amyloid (Aβ) pathology contributes to accelerated brain aging. Using data collected in 1340 cognitively unimpaired participants from 18 to 94 years old selected across multi-site cohorts, we showed that chronological age can be predicted across the whole lifespan from topological properties of graphs constructed from rs-fMRI. We subsequently used the difference between the model-predicted age and the chronological age in pre-symptomatic autosomal dominant AD (ADAD) mutation carriers and asymptomatic individuals at risk of sporadic AD and assessed the influence of genetics and Aβ pathology on brain age. Applying our predictive model in the context of preclinical AD revealed that the pre-symptomatic phase of ADAD is characterized by accelerated functional brain aging. This phenomenon is independent from, and might precede, detectable fibrillar Aβ deposition.

## Introduction

The brain suffers major changes over the course of aging. How the brain ages and how neurodegenerative diseases affect brain aging remains to be fully understood, but increasing evidence suggests that neural systems vulnerable to age are also vulnerable to Alzheimer’s disease (AD) and other age-related neurodegenerative diseases.^1^ Lately, increased availability of large-scale neuroimaging dataset has facilitated the application of machine learning techniques in neuroscience and enabled the development of models that could predict behavior and characteristic traits from brain structure and function, including age.^2–8^ Brain age could be a particularly relevant biomarker in the context of aging and neurodegenerative diseases since,^2^ by predicting age from brain characteristics, it is possible to identify who deviates from the “normal” aging trajectories and which factors influence these deviations. Previous studies found that brain age predictions are influenced by lifestyle factors^9–13^ and other conditions, including AD dementia.^14–16^ In the present study, our main objective was to assess the genetic and pathological factors that could accelerate brain aging in preclinical AD.

AD is defined clinically by a progressive cognitive decline, usually affecting episodic memory first, interfering with daily living activities.^17^ Prior works in AD demonstrated that brain changes occur 2 to 3 decades before symptom onset with the accumulation of cerebral beta-amyloid (Aβ) plaques, followed by the accumulation of tau deposition, functional/metabolic brain alterations, neurodegeneration and, finally, cognitive and functional symptoms.^18,19^ Thus, clinical symptoms are the result of decades-long biological changes and, at the time of diagnosis, alterations are already too massive to be reversed. To better understand disease progression, it is thus crucial to focus on the preclinical phase of AD.^20^ Evaluating individuals in this “silent” phase of the disease is however challenging as deviations from normal aging trajectories are subtle and difficult to assess in this asymptomatic individuals. Autosomal dominant AD (ADAD) is a rare and early onset variant of AD, caused by a mutation on the amyloid precursor protein (*APP*), presenilin 1 (*PSEN1*) or presenilin 2 (*PSEN2*) genes, all involved in Aβ production.^19,21^ These mutations are fully penetrant and the disease progression predictable, making ADAD an ideal model to study the preclinical (i.e. presymptomatic) phase of AD. In sporadic AD (sAD), it is still impossible to determine who will develop dementia with certainty, but some factors have been shown to increase the risk of developing the disease. Amongst them, the apolipoprotein E4 (*APOE*4), known to be involved in Aβ clearance,^21–24^ is the main genetic risk factor of late onset AD. Familial history of sAD has also been associated with a 2- to 4-fold increased incidence of developing AD.^25–29^ Therefore, pre-symptomatic ADAD and asymptomatic individuals with a family history of sAD represent privileged populations to further understand how the disease, in its asymptomatic stage, influence brain aging. Some studies suggested that AD is characterized by an increased structural brain age in symptomatic individuals (mild cognitive impairment [MCI] and AD-demented patients).^14–16^ Our main objective is to test if the disease influences brain aging trajectories early on, prior to dementia.

To assess brain aging in the context of the preclinical phase of AD we aimed first at developing a model able to predict chronological age from resting state functional magnetic resonance imaging (rs-fMRI). Functional brain changes have been suggested to be a sensitive early marker of brain alterations/changes in various conditions, including AD.^30–34^ We used measures of network integration and segregation, known as graph metrics,^35^ as a proxy of global brain functioning. In a second objective, we evaluated how preclinical AD related to brain aging, and more particularly, whether functional brain age differed by genetic factors or Aβ pathology. We included 1624 cognitively unimpaired participants from 18 to 94 years old, recruited and scanned in different studies and centers. We developed a neural net to predict brain age from rs-fMRI. Briefly, we trained the model on cognitively unimpaired individuals ranging in age from 18 to 90 years old and validated its generalisability on a group of cognitively unimpaired individuals from an independent study/site. After validation of the model, we tested if genetic factors or Aβ pathology were associated with accelerated neural aging in “unseen” participants that were not used to develop or validate the brain age model. Pre-symptomatic ADAD mutation carriers were characterized by an accelerated functional brain aging, that was independent from Aβ pathology detected using PET imaging. In an at-risk cohort for sAD, neither the *APOE*4 status nor Aβ pathology was associated with brain aging.

## Results

We gathered rs-fMRI data collected in 1624 cognitively unimpaired participants from 18 to 94 years old, provided by the Dominantly Inherited Alzheimer Network (DIAN), Pre-symptomatic Evaluation of Experimental or Novel Treatments for Alzheimer’s Disease (PREVENT-AD), Cambridge Centre for Ageing and Neuroscience (CamCAN), 1000-Functional Connectomes Project – Cambridge site (FCP-Cambridge), Alzheimer’s disease Neuroimaging Initiative (ADNI) and International Consortium for Brain Mapping (ICBM) cohorts, to build a “brain age” predictive model. A harmonized preprocessing pipeline was applied to all individuals and 26 graph metrics, chosen based on their ability to quantify whole-brain connectivity, were extracted from each participant’s correlation matrix (see Material and Methods for details).

After processing and quality control, 1340 individuals remained for the analyses and were divided into training, validation, and test sets as detailed in Table 1. More specifically, the data set was divided such that the training and validation sets were representative of healthy aging. Mutation non-carriers from DIAN (~75% of DIAN non-carriers, randomly selected) and a few individuals from the PREVENT were assigned to the training set, along with individuals from the FCP-Cambridge, and part of the cognitively unimpaired individuals selected randomly from the CamCAN and ADNI. ICBM was used as an independent sample of healthy individuals, which despite being of modest size covers the entire adulthood, to assess the generalisability of the brain age model to other datasets (validation set). Finally, the test set included our population of interest (DIAN mutation carriers, most PREVENT-AD participants) and the remaining asymptomatic individuals from the other cohorts (DIAN mutation non-carriers, CamCAN and ADNI participants). Importantly, each individual was included in only one of the three sets such that no overlap exists between the training, validation and test sets (see Methods for details). Furthermore, the analyses were only performed on the test set after the brain age model was fully developed and validated.

**Table 1.** Data sets characteristics (sample size [age range in years])

| **Cohorts** | **Training set** | **Validation set** | **Test set** |
| --- | --- | --- | --- |
| **DIAN mutation non-carriers** | 105<br>[19-69 yo] | - | 29<br>[20-58 yo] |
| **DIAN mutation carriers** | - | - | 125<br>[18-61 yo] |
| **PREVENT-AD** | 36<br>[55-78 yo] | - | 256<br>[55-84 yo] |
| **CamCAN** | 408<br>[18-87 yo] | - | 96<br>[18-88 yo] |
| **ADNI** | 29<br>[66-94 yo] | - | 15<br>[65-90 yo] |
| **FCP-Cambridge** | 195<br>[18-30] | - | - |
| **ICBM** | - | 46<br>[19-79 yo] | - |
| **Total sample** | **773**<br>**[18-94 yo]** | **46**<br>**[19-79 yo]** | **521**<br>**[18-90 yo]** |

### Brain age model

First, in order to reduce the number of inputs to the model, we searched for the graph metrics that were the most reliably predictive of chronological age.^4^ To do so, the training set data was entered in parallel to a support vector machine (SVM) and a regression tree ensemble models to estimate which graph metrics were the most important to predict chronological age (*i.e.*, highest weights). Graph metrics were then ranked separately by order of SVM weights and ensemble model importance (*i.e.*, highest load corresponding to the most important). The average rank from both models was used to determine the overall importance of each metric presented in Figure 1A).

**Figure 1.**
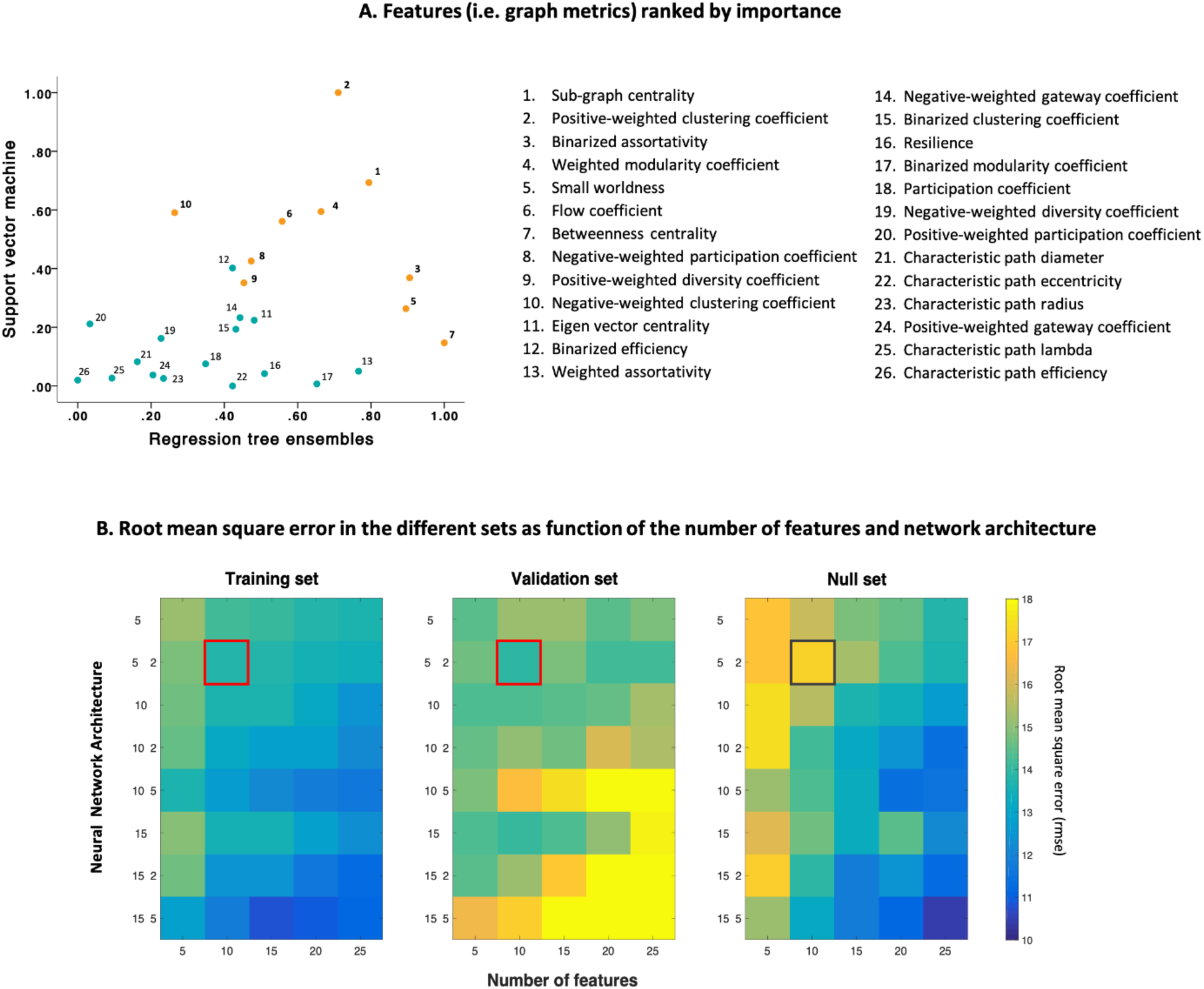
Scatter plots (A) of SVM model weights (y-axis) and decision tree feature importance (x-axis). Model weights are absolute value, and normalized such that 1 indicates highest importance. Numbers next to data points indicate their rank (i.e. 1 = highest average rank between both SVM and ensemble models). Root mean square error (rmse) of different neural network models (B) with inputs sorted according to rank for the training set (left), and the validation set (middle). Neural networks trained with randomly-ranked inputs served as our null models (right). The x-axis indicates the number of inputs into the model (number of graph metrics) while the y-axis indicates the network architecture. For example, ‘5’ means 1 hidden layer with 5 units, ‘5 2’ means 2 hidden layers, the first one with 5 units and the second with 2 units. Darker colors indicate higher accuracy. The red square identifying the model that provides the better generalisability in the validation set (lowest rmse) contains 2 hidden layers of 5 and 2 units, and uses the 10 highest-ranked graph metrics as input. The same neural network trained on randomly-ranked inputs (null model, grey square) provides lower accuracy.

In a second step, we created an optimized model requiring the fewest number of features possible. The neural network was optimized by i) generating different models using the training set and ii) evaluating which one provided the better generalisability to the validation set (*i.e.*, avoid overfitting and give the better prediction on an independent set). Overfitting happens when a model fits too well the data and noise of a specific dataset, making generalisability of the model to other data unlikely. The validation set helped us to determine the best trade-off between optimizing the age prediction in the training set and enabling a good generalisability. In addition, to ensure the relevance of metrics ranking conducted in the first step, performance of neural nets on the training data, when accounting for this feature ranking, was also compared to the performance of the same neural nets while including features randomly (*i.e., “*null model”). Root mean square error (rmse) was used to assess the model performance and, to allow for a more precise estimation in this selection process, the averaged rmse over 3 iterations was considered for each model. Different neural networks were built, from simpler to more complex, varying in number of input features (5, 10, 15, 20, or 25 more important graph metrics, according to the ranking determined previously), hidden layers, and hidden layer units.

In general, increasing model complexity (more features and hidden layers/units) lead to better performance in the training set (Figure 1B, left panel). However, and as expected, too much complexity resulted in overfitting; while performance increased as the model complexity increased on the training set, performance decreased in the validation set, suggesting that simpler models provide better generalisability (Figure 1B, middle panel). In the null model, the same rationale was adopted on training data, but graph metrics were entered progressively in a random order (see Figure 1B, right panel). This null model indicated that the neural network performed better when features were ordered by SVM weights and ensemble feature importance (as determined previously), at least with the simpler models.

The model that allowed the lowest rmse in the validation set (averaged rmse over 3 iterations = 13.89) had 10 inputs (i.e., the 10 most important metrics, see Figure 1A) and 2 hidden layers (5 units in the first layer and 2 in the second). The performance from this specific model was similar to the one obtained on the training set (averaged rmse over 3 iterations = 13.75). This model was thus applied to the remaining unseen data (test set) to test if genetics or AD pathology accelerate functional brain aging.

The association between chronological age and the predicted age provided by the model is depicted for each dataset in Figure 2. As expected, the predicted age was correlated with chronological age in the training set (R^2^=.53, p < .0001; rmse=14.01, mean absolute error [mae] = 11.00; Figure 2A) and the validation set (R^2^=.49, p < .0001; rmse=13.84; mae = 11.90; Figure 2B). The model was also able to predict chronological age from functional brain properties in the test set (R^2^=.36, p < .0001; rmse=13.24; mae = 11.58; Figure 2C).

**Figure 2.**
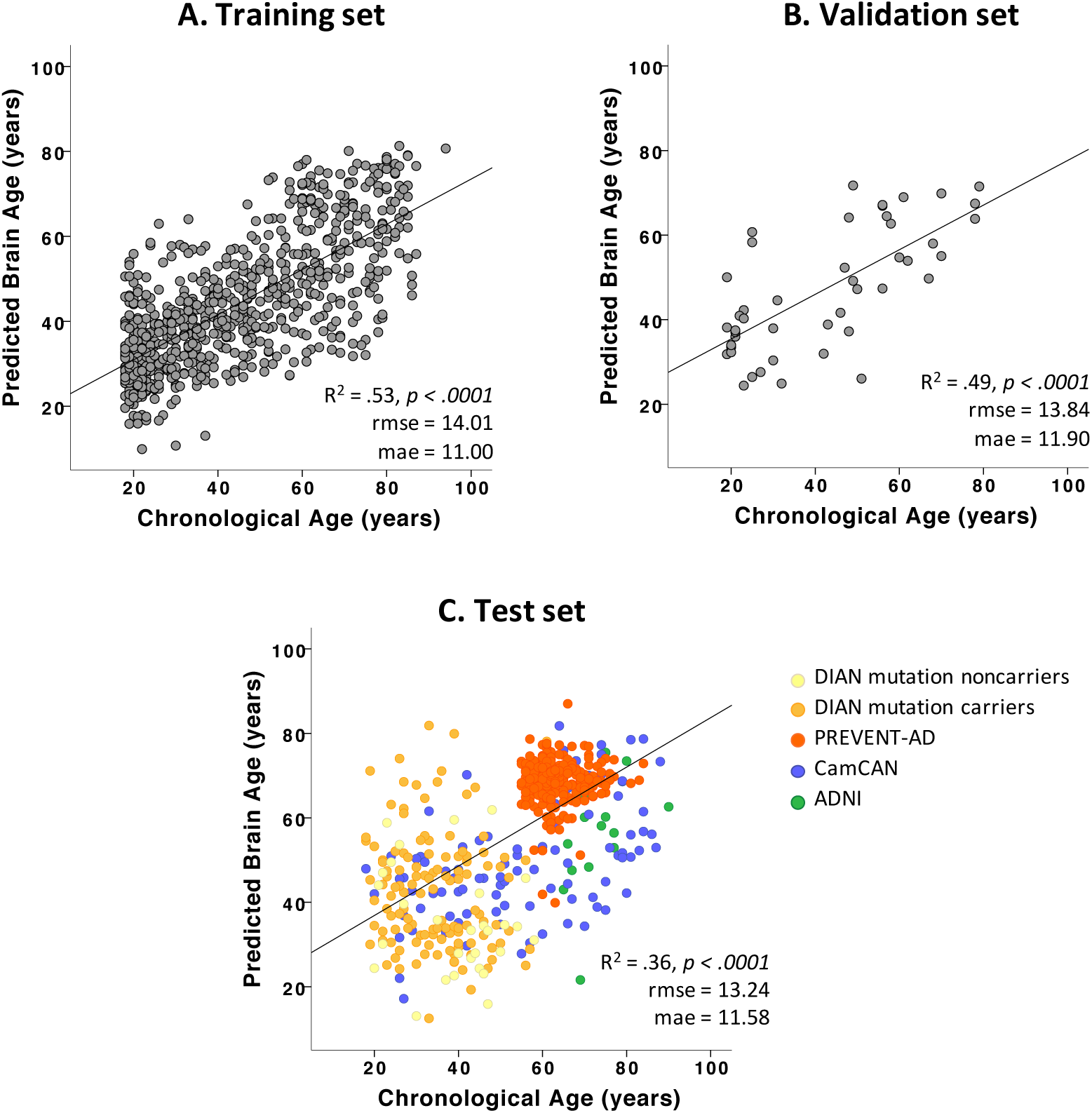
Brain Age model performance across data sets. Correlations between chronological age (x-axis) and age predicted by the neural network (y-axis) are represented for the training (A), validation (B) and test (C) sets. rmse: root mean square error; mae: mean absolute error

In order to assess the characteristics of functional brain aging in preclinical AD and evaluate whether genetic determinism/risk and Aβ pathology were related to accelerated brain aging, we calculated the predicted age difference [PAD = predicted – chronological age]^3^ for each participant in the test set and included this measure in further analyses.

### Functional brain aging in the asymptomatic phase of Alzheimer’s disease

Analyses were conducted on the DIAN and PREVENT-AD participants included in the test set (Table 2).

**Table 2.**
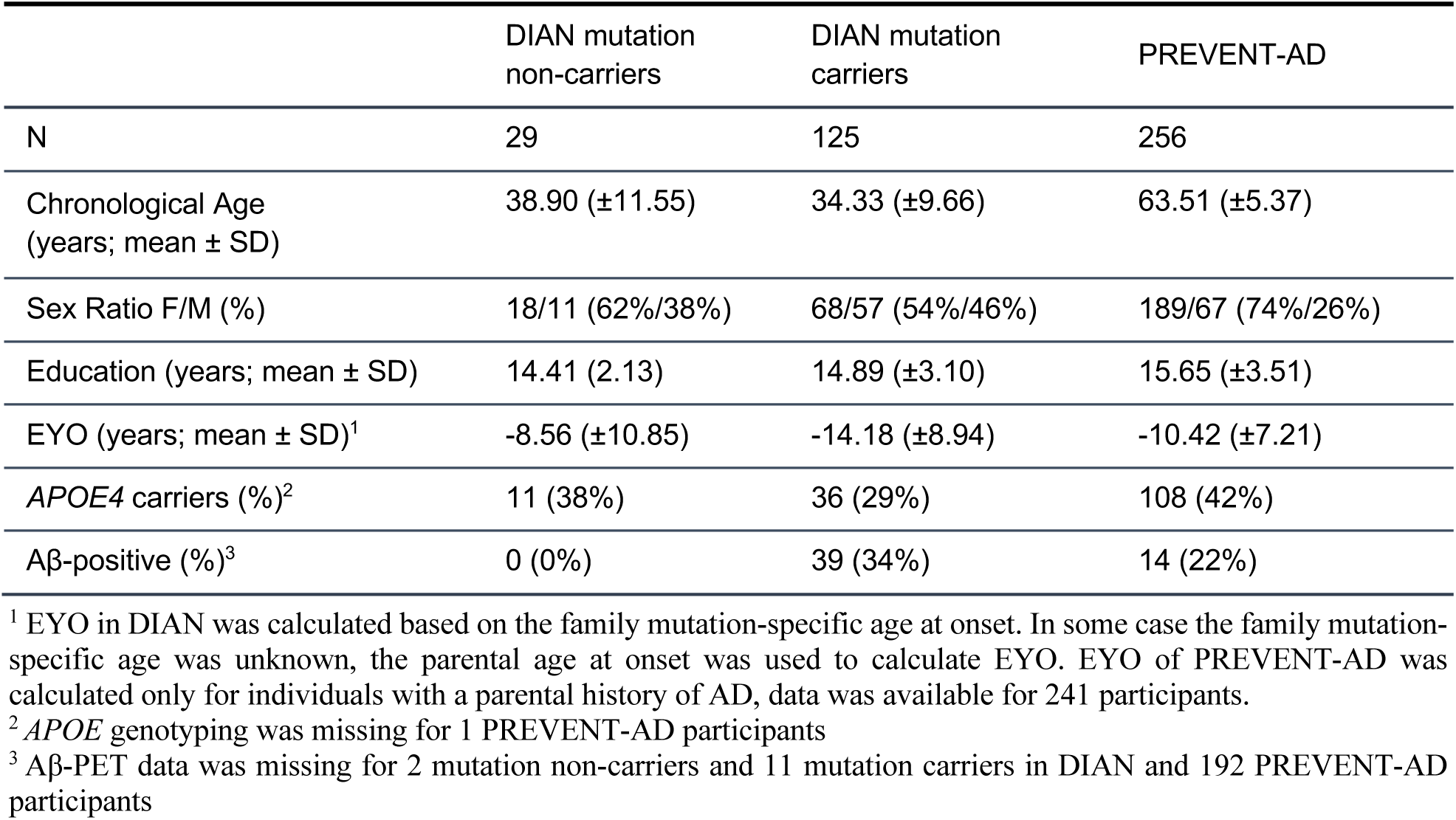
DIAN and PREVENT-AD test set characteristics

|  | DIAN mutation non-carriers | DIAN mutation carriers | PREVENT-AD |
| --- | --- | --- | --- |
| N | 29 | 125 | 256 |
| Chronological Age (years; mean ± SD) | 38.90 ($\pm 11.55$) | 34.33 ($\pm 9.66$) | 63.51 ($\pm 5.37$) |
| Sex Ratio F/M (%) | 18/11 (62%/38%) | 68/57 (54%/46%) | 189/67 (74%/26%) |
| Education (years; mean ± SD) | 14.41 (2.13) | 14.89 ($\pm 3.10$) | 15.65 ($\pm 3.51$) |
| EYO (years; mean ± SD)<sup>1</sup> | -8.56 ($\pm 10.85$) | -14.18 ($\pm 8.94$) | -10.42 ($\pm 7.21$) |
| APOE4 carriers (%)<sup>2</sup> | 11 (38%) | 36 (29%) | 108 (42%) |
| Aβ-positive (%)<sup>3</sup> | 0 (0%) | 39 (34%) | 14 (22%) |

<sup>1</sup> EYO in DIAN was calculated based on the family mutation-specific age at onset. In some case the family mutation-specific age was unknown, the parental age at onset was used to calculate EYO. EYO of PREVENT-AD was calculated only for individuals with a parental history of AD, data was available for 241 participants.
<sup>2</sup> APOE genotyping was missing for 1 PREVENT-AD participants
<sup>3</sup> Aβ-PET data was missing for 2 mutation non-carriers and 11 mutation carriers in DIAN and 192 PREVENT-AD participants

To test if genes involved in AD, which are either responsible of ADAD or increase the risk of sAD, were associated with accelerated brain aging, we compared PAD between mutation non-carriers *vs* mutation carriers from DIAN, and *APOE*4 carriers *vs* non-carriers in the PREVENT-AD. These analyses were only performed in the test set. Model’s prediction in DIAN mutation carriers overestimated their chronological age (*i.e.*, positive PAD = 8.19 years), and this PAD was higher than it was in the mutation non-carriers (F_1,152_=10.12; p=.002; Table 3 & Figure 3A1), in whom the model’s prediction underestimated the chronological age (PAD = −3.54 years). Overall, the predicted age in the PREVENT-AD cohort overestimated the chronological age by ~5 years, but *APOE*4 status was not associated with this prediction (F_1,253_<1; p=.69, Table 3 & Figure 3B1).

**Table 3.** Model’s prediction according to the presence of genetic mutation/risk and Aβ pathology in DIAN and PREVENT-AD cohorts (test set only).

|  | DIAN |  |  |  | PREVENT-AD |  |  |  |
| --- | --- | --- | --- | --- | --- | --- | --- | --- |
| | Mutation<br>NonCarr.<br>Aβ- | Mutation<br>Carriers<br>all | Mutation<br>Carriers<br>Aβ- | Mutation<br>Carriers<br>Aβ+ | *APOE4*<br>NonCarr. | *APOE4*<br>Carriers | Aβ- | Aβ+ |
| N | 29<sup>1</sup> | 125 | 75 | 39 | 147 | 108 | 50 | 14 |
| Chronological<br>Age in years<br>(±SD) | 38.90<br>(±11.55) | 34.35<br>(±9.66) | 33.15<br>(±9.09) | 38.18<br>(±10.04) | 63.89<br>(±5.74) | 63.05<br>(±4.77) | 62.90<br>(±4.28) | 64.71<br>(±5.69) |
| Predicted Age<br>in years (±SD) | 35.35<br>(±12.90) | 42.51<br>(±13.90) | 41.55<br>(±13.19) | 41.18<br>(±14.09) | 68.94<br>(±5.14) | 68.46<br>(±5.01) | 68.80<br>(±6.08) | 70.37<br>(±2.97) |
| PAD in years<br>(±SD) | -3.54<br>(±18.99) | 8.19<br>(±17.63) | 8.40<br>(±17.21) | 6.62<br>(±18.09) | 5.04<br>(±7.62) | 5.41<br>(±6.79) | 5.90<br>(±7.60) | 5.66<br>(±6.56) |

Aβ - = amyloid-negative; Aβ + = amyloid-positive; Nonarr. = Non-Carriers; PAD = Predicted age difference; SD = standard deviation
<sup>1</sup> Aβ-PET data was missing for 2 mutation non-carriers. To avoid further reducing the number of individuals in this group, they were both exceptionally kept for the corresponding analysis and their Aβ negativity was determined based on i) their CSF data (available for both of them) and ii) the unlikelihood that they have detectable Aβ pathology on PET considering their young age and the absence of ADAD mutation. Importantly, results remained the same when removing those two individuals from the analysis (data not shown).

**Figure 3.**
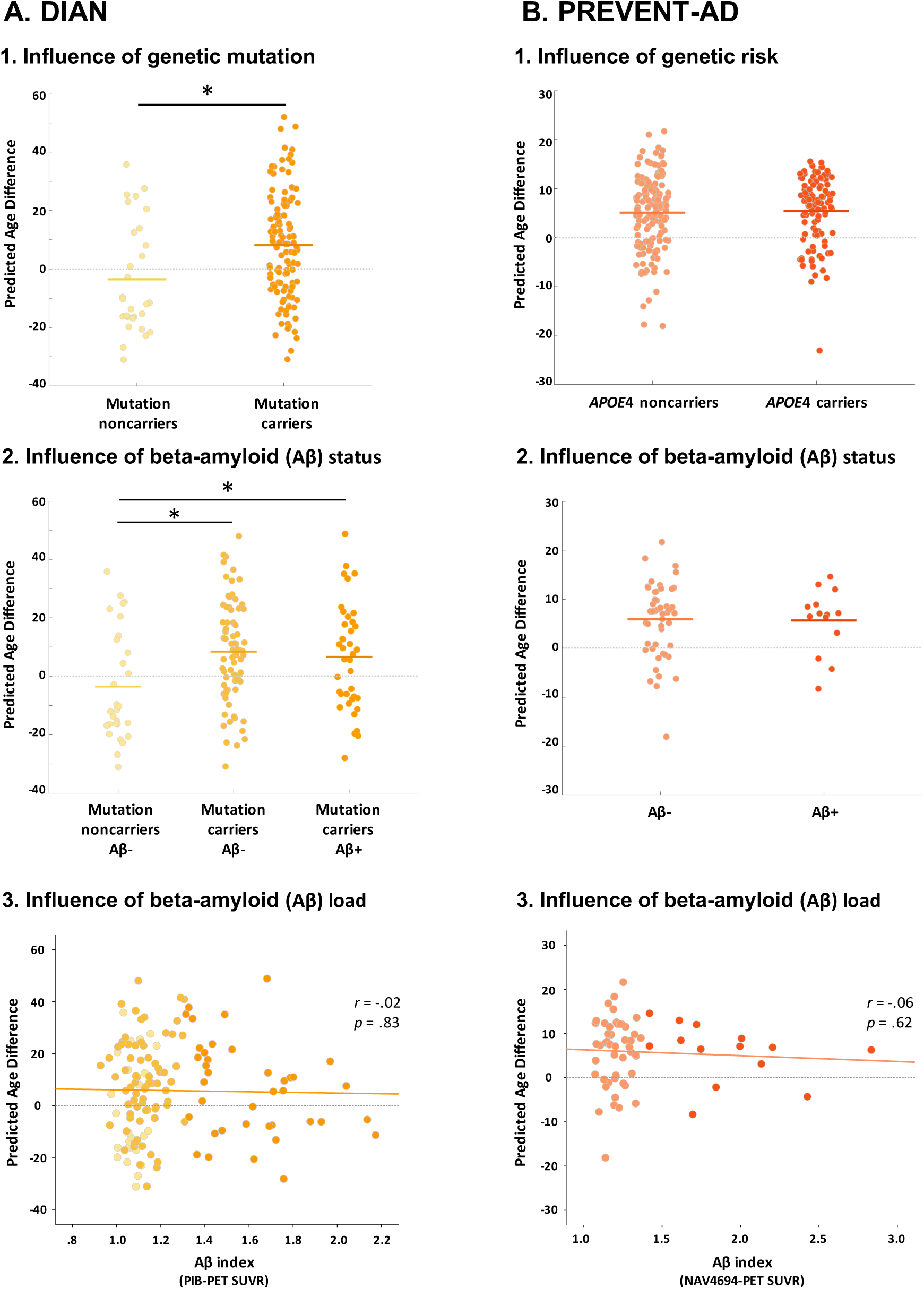
Difference between the age predicted from brain function and actual age (PAD) in DIAN (A) and PREVENT-AD (B). Brain age is overestimated in autosomal dominant mutation carriers compared to non-carriers (A.1.) but this overestimation in mutation carriers is not associated with beta-amyloid (Aβ) status (A.2) or load (A.3). In individuals at risk of sporadic Alzheimer’s disease, brain age is overestimated irrespectively of *APOE4* genotype (B.1.) and Aβ status (B.2) or load (B.3). Of note, Aβ-PET acquisition and processing differed between DIAN and PREVENT-AD, preventing direct comparison of their SUVR range.

Considering the importance of Aβ deposition in the cascade of event leading to AD, we assessed how Aβ burden related to on functional brain aging. To do so we assessed the effect of Aβ deposition, measured by PET, on the PAD in both the DIAN and PREVENT-AD cohorts. Aβ-PET was acquired using C^11^-PIB in DIAN and F^18^-NAV4694 in PREVENT-AD, and Aβ burden was determined for each cohort according to their own processing pipelines and methods (see Methods section for Aβ measurements details). Influence of Aβ burden on functional brain aging was investigated both by comparing Aβ+ and Aβ-individuals (dichotomous variable), and by assessing the correlation between PAD and Aβ load (continuous variable). When applicable, post-hoc comparisons were done using Fisher’s LSD. Amongst DIAN test set’s participants, all mutation non-carriers were Aβ-negative (n= 29), 75 mutation carriers were Aβ-negative and the remaining 39 carriers were Aβ-positive. The PAD varied across the three groups (F_1,152_=4.81; p=.01 Table 3 & Figure 3A2). Post-hoc comparison showed that the PAD did not differ between Aβ-negative and Aβ-positive mutation carriers (mean PAD difference between Aβ-negative and Aβ-positive carriers = 1.78 years; p=.87; Table 3), but both groups differ from the Aβ-negative mutation non-carriers (mean PAD difference between Aβ-negative non-carriers and Aβ-negative carriers = −11.94 years, p=.003; and mean PAD difference between Aβ-negative non-carriers and Aβ-positive carriers = −10.17 years, p=.021; Table 3). Using the Aβ burden index as a continuous variable, there was no association between Aβ load and the PAD (r_143_=-.02, p=.83; Figure 3A3). Reducing the analysis to mutation carriers only provided similar results (r_114_=-.12, p=.61). These results suggest that the acceleration of brain age in the mutation carriers was not associated with Aβ pathology as measured by PET, and was present even in individuals that do not have detectable Aβ pathology yet.

In PREVENT-AD, amongst the 64 individuals who underwent Aβ-PET imaging (test set only), 50 were classified as being Aβ-negative and 14 were Aβ-positive. The PAD was not associated with Aβ burden, either when looking at the influence of Aβ-positivity (F_1,62_<1; p=.92, Table3 & Figure 3B2) or looking at Aβ load (r_64_=-.06; p=.62; Figure 3B3). Adding the delay between PET and rs-fMRI assessments as a covariate provide similar results (F_1,61_<1; p=.84 using Aβ-positivity and r_61_=-.08; p=.55 for the partial correlation with Aβ load).

## Discussion

Machine learning is becoming more and more popular in the field of neuroscience. Combined with larger and more available datasets, these techniques will improve our understanding of brain function, and our ability to predict health trajectories from brain properties.^1^ Previous models of brain age have been informed primarily by characteristics of brain structure.^16.^ Changes in brain function, however, could be a more sensitive and earlier marker of brain abnormalities, notably in conditions in which structural changes are not prominent or occur late in the disease progression. Brain functional changes have been hypothesised to be among the earliest brain changes related to AD.^30,34,36,37^ Here we developed a model able to predict brain age across the whole adulthood (18-94) using topological properties of graphs, constructed from rs-fMRI.

The most common ways of approaching functional connectivity in aging and AD have involved the assessment of some specific networks’ hyper- or hypo-connectivity. The large existing literature on the subject suggests connectivity disruptions through the course of these two processes, notably targeting the default mode network.^38–45^ We took advantage of global brain function, while applying graph metrics,^7,35^ to assess information integration in the brain. This approach provides a holistic view of brain function as it relates to age and disease proclivity and has been shown previously to change through aging and AD.^46^

Our study indicates that it is possible to predict brain age from rs-fMRI, taking advantage of global measures of network integration and segregation.^7,35^ Our proposed model had been developed on participants from different cohorts and sites and validated on an independent monocentric dataset to ensure generalisability of our model to other data sets. While being of modest size, this validation set represents a completely independent dataset that covers the entire adulthood. Importantly, the test set was never used in the development/validation of the brain aging model. Also, the model was not tuned any further after the hypotheses were tested meaning that the hypotheses were only tested once on a model that we believe was optimal. This ensured that our results regarding brain aging in the context of pre-symptomatic AD were independent from how the model was built. Facing the potential problems of an aging population, researchers from different fields are now trying to further understand aging and predict early deviation from healthy aging trajectories, with the final objective to prevent neurodegenerative diseases. A generalisable model of brain aging, as developed here, could thus be quite valuable in future aging studies.

Applying our model in the context of AD, we found evidence of accelerated brain functional aging in individuals in the pre-symptomatic phase of ADAD, independently from the amount of Aβ pathology measured in PET. Studies of ADAD have shown that biomarkers such as CSF-Aβ, start changing in mutation carriers as early as 25 years before symptom onset. This is followed by the accumulation of Aβ deposition objectified by PET imaging (~15 years before symptom onset), changes in tau concentrations measured by CSF and cerebral atrophy (~15 years before onset), glucose hypometabolism and episodic memory decline (~10 years before onset), and global cognitive decline (~5 years before onset).^19,47^ One major hypothesis in the field of AD is that Aβ pathology triggers the cascade of biomarkers changes that will lead to the clinical expression of the disease.^48^ This seems particularly true in the case of ADAD, which is due to the mutation of a gene involved in Aβ production. Here, we found that the brain age of ADAD mutation carriers was overestimated when compared to non-carriers. This suggests that the pre-symptomatic phase of AD is accompanied by an accelerated brain aging. More specifically, this effect was not associated with Aβ burden and was already observed in mutation carriers who did not reach Aβ-positivity, suggesting that accelerated functional brain aging occurs before Aβ accumulation can be detected using PET imaging. While we cannot exclude the fact that some Aβ negative individuals would in fact be Aβ accumulators^49–51^ or present other forms of Aβ that cannot be detected through PET, this could also suggest that mutated genes have life-long effects on the brain that are not fully dependent on Aβ accumulation. Consistently, a previous study in *PSEN1* mutation carriers from the Columbian cohort showed early changes in brain function before evidence of cerebral Aβ plaque accumulation.^52^ This also aligns with studies in young *APOE*4 carriers that showed brain structure and function changes associated with the presence of the e4 allele in young adults, and even in children.^53–57^ While *APOE*4 has often been suggested to increase the risk of sAD through its role in Aβ accumulation,^22,58–62^ the existence of an effect of *APOE*4 in early life (*i.e.* before evidence of Aβ accumulation) indicates that *APOE*4 also act on the brain integrity through Aβ-independent pathways.^24^ Altogether, this suggests that AD genetic mutations can influence brain properties early in life, independently from fibrillar Aβ deposition, and that functional brain changes might precede this phenomenon,^see also 36^ at least in ADAD.

In individuals with a family history of sAD (PREVENT-AD), functional brain age was found to be overestimated, while it was not the case in other site/cohorts of similar age-range. While it is tempting to interpret these results as being the consequence of their family risk, we believe that this is largely due to a site effect. Beyond all being scanned in the same site and scanner, individuals from this cohort were selected according to the same and very strict inclusion criteria. To minimize the possibility of site effects, we included data from a variety of cohorts and sites, and validated the model on a completely independent validation set (new site). Despite this effort we cannot exclude the possibility that the age overestimation of most PREVENT-AD participants (Figure 2C) could be the result of a site effect. While this phenomenon limits our ability to draw direct comparison between the cohorts, it does not influence our main findings, which are the results of within-cohort comparisons. When investigating the characteristics of PAD within the PREVENT-AD, we did not find a difference between *APOE4* carriers and non-carriers. While this is in contradiction with previous studies suggesting early functional alteration in *APOE*4 carriers (see above), it is possible that the influence of *APOE*4 was not strong enough to influence functional brain aging in this particular sample that is already at increased risk of sAD. Further studies are needed to better understand this result. Furthermore, the absence of association between Aβ burden and the overestimation of brain age found in PREVENT-AD is in line with what was found in DIAN, suggesting that functional brain aging and Aβ accumulation are, in part, occurring independently.^see also 63^

Using rs-fMRI graph metrics, we’ve developed a model that can predict brain age across the whole lifespan. Applying this model to predict brain aging in the context of preclinical AD revealed that the pre-symptomatic phase of ADAD is characterized by accelerated functional brain aging. This phenomenon is independent from, and might therefore precede, fibrillar Aβ accumulation.

## Methods

### Cohorts and participants

#### Dominantly Inherited Alzheimer Network - DIAN

DIAN is a multisite longitudinal study fully described elsewhere,^64^ which enrolls individuals age 18 and older who have a biological parent that carry a genetic mutation responsible for ADAD. They all undergo clinical and cognitive assessments, genetic testing and imaging (magnetic resonance imaging [MRI] and amyloid-positron emission tomography [PET]). Data has been obtained after request and IRB approval (information can be found at dian.wustl.edu/our-research/observational-study/). Baseline data from the cognitively unimpaired mutation carriers and non-carriers archived in the DIAN data-freeze 10 (January 2009 to May 2016) were used in the present study. Baseline data from 280 cognitively unimpaired individuals (mutation carriers and non-carriers) aged between 18 and 69 years old, for whom structural MRI and rs-fMRI data were available, have been included.

#### Pre-symptomatic Evaluation of Experimental or Novel Treatments for Alzheimer’s Disease-PREVENT-AD

The PREVENT-AD (Douglas Mental Health Institute, Montréal) is a monocentric longitudinal cohort, fully described elsewhere.^65^ Briefly, 399 cognitively unimpaired older individuals with a family history of sAD (at least one parent and/or multiple siblings) were enrolled between September 2011 and November 2017. Inclusion criteria included i) being 60 or older; 55 to 59 for individuals who were less than 15 years from the age of their relatives at symptom onset, ii) being cognitively normal and iii) no history of major neurological or psychiatric disease. Participants underwent clinical and cognitive examinations, blood tests and MRI annually. Data from the present study were archived in the data release 5 and are partially available at https://openpreventad.loris.ca/. PET scans were acquired in a subset of participants between February 2017 and July 2019. Three hundred and fifty-three participants, aged 55 to 84, for whom baseline structural MRI and rs-fMRI were available were included in the present study.

#### Cambridge Centre for Ageing and Neuroscience - CamCAN

The Cambridge Centre for Ageing and Neuroscience (Cam-CAN; http://www.cam-can.org/) is a large-scale collaborative research project, launched in October 2010, using epidemiological, behavioural, and neuroimaging data to characterise age-related changes in cognition and brain structure and function, and to uncover the neurocognitive mechanisms that support healthy cognitive ageing.^66,67^ In the present study, 648 individuals aged between 18 and 88, with structural MRI and rs-fMRI data were included.

#### Alzheimer’s disease Neuroimaging Initiative - ADNI

ADNI data used in the preparation of this article were obtained from the Alzheimer’s Disease Neuroimaging Initiative (ADNI) database (adni.loni.usc.edu).^68,69^ The ADNI was launched in 2003 as a public-private partnership, led by Principal Investigator Michael W. Weiner, MD. The primary goal of ADNI has been to test whether serial MRI, PET, other biological markers, and clinical and neuropsychological assessment can be combined to measure the progression MCI and early AD. Forty-nine cognitively unimpaired individuals with structural MRI and rs-fMRI data were included in the present study.

#### 1000-Functional Connectomes Project (Cambridge) – FCP-Cambridge

The 1000-Functional connectomes project (FCP) is a large initiative that gathers functional data from cognitively unimpaired adults recruited worldwide (33 sites) and makes it publicly available to facilitate discovery science of brain function (http://fcon_1000.projects.nitrc.org/fcpClassic/FcpTable.html).^38^ We used the large dataset from Cambridge-Buckner that includes 198 subjects between 18-30 years old collected at the Cambridge site ([FCP-Cambridge], PI: Buckner, R.L.).

#### International Consortium for Brain Mapping - ICBM

The ICBM dataset^70^ is publicly available as part of the 1000-FCP repository (see above; see also ^71^ for details). The dataset is constituted of 86 cognitively unimpaired older adults from 19 to 95 years old who underwent structural MRI and rs-fMRI at the same site (Montreal Neurological Institute, Canada).

For the purpose of the brain age model, participants were divided into training, validation, and test sets. To build a model of “healthy” brain aging, mutation non-carriers from DIAN and a few individuals with limited genetic risk from the PREVENT AD (selected based on their *APOE4* genotype) were assigned to the training set, along with individuals from the FCP-Cambridge and part of the cognitively unimpaired individuals selected randomly from the CamCAN and ADNI (training set). ICBM was used as an independent sample of healthy individuals to assess the generalisability of the brain age model to other datasets (validation set). Finally, the test set included our population of interest (DIAN mutation carriers, PREVENT-AD participants) and the remaining asymptomatic individuals from the other cohorts (DIAN mutation non-carriers, CamCAN and ADNI participants).

### Standard Protocol Approvals, Registrations, and Participants Consents

All studies were approved by study sites’ respective regional ethics committees. All participants gave written informed consent prior to participation.

### Data availability

All data used in the present study are either publicly available (PREVENT-AD MRIs; CamCAN; 1000-FCP datasets) or can be shared upon reasonable request and approval by the study scientific committees and/or institutional review boards (DIAN; PREVENT-AD additional measurements, including PET; ADNI).

### MRI acquisition and processing

#### DIAN

DIAN imaging data was acquired at multiple sites on 3T scanners by applying ADNI parameters and procedures.^19,47,64^ T1-weighted MRI (used for rs-fMRI processing) were acquired with the following parameters: repetition time (TR) = 2400ms, echo time (TE) = 16ms, flip angle = 8°, acquisition matrix = 256×256, voxel size = 1×1×1mm. Eyes-open rs-fMRI images were acquired using the following parameters: TR = 2230ms or 3000ms; TE=30 ms, flip angle=80°, voxel-size = 3.3×3.3×3.3mm, field of view (FOV) = 212, 140 volumes; acquisition lasting approximately 5min or 7min.

#### PREVENT-AD

MRI data were acquired on a 3T Magnetom Tim Trio (Siemens) scanner. T1-weighted images were obtained using a GRE sequence with the following parameters: TR = 2300ms; TE = 2.98ms; flip angle = 9°; matrix size = 256×256; voxel size = 1×1×1mm; 176 slices. For resting state fMRI scans, two consecutive functional T2*-weighted scans were collected eyes-closed with a blood oxygenation level-dependent (BOLD) sensitive, single-shot echo planar sequence with the following parameters: TR = 2000ms; TE = 30 ms; flip angle = 90°; matrix size = 64×64; voxel size = 4×4×4 mm; 32 slices; 150 volumes, acquisition time = 5min45s. For consistency with the other cohorts that only had one run, only the first run was considered for each participant.

#### CamCAN

Images were acquired on a 3T Magnetom Tim Trio syngo (Siemens). T1-weighted MRI were acquired using the following parameters: 3D MPRAGE GRAPPA=2, TR = 2250ms, TE = 2.99ms, TI = 900ms; flip angle = 9°; voxel-size 1mm isotropic; FOV = 256×240×192mm; acquisition time = 4mins 32s. Rs-fMRI data were acquired eyes closed using a T2* GE EPI sequence with the following parameters: TR = 1970ms; TE = 30ms, flip angle = 78°; voxel-size = 3×3×4.44mm, FOV = 192×192; 261 volumes of 32 axial slices 3.7mm thick with a 0.74mm gap, acquisition time = 8mins 40s.

#### ADNI

Data were acquired on multiple sites, following the ADNI protocol.^68,72^ Structural images were acquired using a 3D MPRAGE T1-weighted sequence with the following parameters: TR = 2300ms; TE = 2.98ms; TI = 900ms; flip angle = 9°; voxel size=1.1×1.1×1.2mm^3^; FOV = 256×240 mm^2^; 170 slices. The rs-fMRI images were obtained, eyes open, using a T2 weighted echo-planar imaging sequence with the following parameters: TR = 3000ms; TE = 30ms; flip angle = 80°; 48 slices of 3.3mm; 140 volumes; acquisition lasting approximately 5minutes.

#### FCP-Cambridge

Images were acquired using a Siemens 3 T Trio scanner. High-resolution T1-weighted images were acquired as follows: MP-RAGE TR = 2200ms, TE = 1.04–7.01ms, flip angle = 7°, voxel size = 1.2×1.2×1.2mm, FOV = 230mm, 144 sagittal slices. Rs-fMRI were collected, eyes open, with the following parameters: EPI TR = 3000ms, TE = 30ms, flip angle = 85°, voxel size=3×3×3mm, FOV = 216mm, 47 axial slices, 124 volumes, lasting approximately 6 minutes.

*ICBM* data was acquired on a Siemens Sonata 1.5T MR scanner at the MNI. T1-weighted scan was acquired as follows: TR = 2200ms, TE = 92ms, flip angle = 30°, 256×256 matrix with a 1×1 mm^2^ resolution, 176 contiguous sagittal slices covering the whole brain, slice thickness = 1 mm. Three rs-fMRI runs were acquired eyes closed with the following parameters: 2D echoplanar BOLD MOSAIC sequence, TR = 2000ms, TE = 50ms, flip angle= 90°, 64×64 matrix with a 4×4mm^2^ resolution, 23 contiguous axial slices covering the cortex but not the cerebellum, slice thickness = 4mm, 138 volumes; lasting approximately 4min30.. For consistency with the other cohorts that only had one run, only the first run was considered for each participant.

### Rs-fMRI processing

All functional images were preprocessed the same way using the NeuroImaging Analysis Kit version 0.12.4 (NIAK; http://niak.simexp-lab.org/), as described in previous publications.^73,74^ Briefly, images underwent slice timing correction, and rigid-body motion parameters were estimated. T1-weighted images were linearly and non-linearly normalized to the MNI space. After coregistration to structural scans, functional images were normalized to the MNI space by applying parameters from the T1-weighted images and resampled to 2mm isotropic. Slow time drifts, average white matter and cerebrospinal fluid signal and motion artifacts (first principal components of the six realignment parameters, and their squares) were regressed out from the rs-fMRI time series. Finally, fMRI volumes were smoothed with a 6mm Gaussian kernel. Frame displacement was calculated and those exhibiting displacement >0.5 were removed (scrubbed), along with one adjacent frame prior, and two consecutive frames after.^75^ Images with less than 40% of their original data after scrubbing were discarded. Overall, 260 individuals (16 DIAN, 60 PREVENT-AD, 130 CamCAN, 14 ADNI, 1 FCP-Cambridge and 39 ICBM) were discarded due to failing preprocessing standards or having insufficient data after scrubbing.

Average BOLD signals was extracted from 272 regions corresponding to the Power and Petersen functional atlas,^76^ to which key regions of the limbic system were added.^77^ Regions labeled as ‘uncertain’, or with weak or non-existent signal in any one image were excluded from all images, resulting in 238 total regions (see supplement eTable-1 for the total listing of the regions). For each subject, BOLD activity time series from these regions were used to construct a 238×238 Pearson correlation matrix, which was then Fisher’s Z-transformed.

Motion-related noise was further mitigated using the mean regression (‘MR’) technique as outlined previously.^78^ Briefly, the average of all correlation values within the upper diagonal of the correlation matrix was calculated for each subject in the training data. A linear fit between these across-subject average values and the across-subject value at each element of the correlation matrix was generated, creating a slope and intercept term associated with each element of the matrix. The final value used in each element of the correlation matrix was equal to the residual between the MR-model fit and the original correlation value. Importantly, the MR model was created with only the training data.

For each subject, 26 graph metrics, chosen based on their ability to quantify whole-brain connectivity, were extracted from the correlation matrix using the Brain Connectivity Toolbox (https://sites.google.com/site/bctnet/),^35^ in Matlab. Both weighted and unweighted metrics were calculated, if applicable. In the case of unweighted metrics, correlation matrices were thresholded at 5% link density, which ensured only the top 5% strongest correlation values were counted as connections in the matrix.^79^ One global value was extracted for each graph metric. In cases where a metric was outputted for each region (e.g. subgraph centrality), the median or median of log values was used as a global estimate. Small-worldness and resilience metrics, not included in the toolbox but both shown to be strong indicators of age, were calculated as previously described.^7^ Subjects with any graph metric that was 5 standard deviations beyond the training set group mean was removed from the analysis entirely. A total of 15 individuals from the training set (1 DIAN mutation noncarrier, 11 CamCAN, 2 FCP-Cambridge, 1 ADNI), 1 from the validation set (ICBM) and 8 from the test set (1 DIAN mutation non-carriers, 3 DIAN mutation carriers, 1 PREVENT-AD, 3 CamCAN) were excluded.

### Brain age model

The general procedure for iterating through different models included 5-fold cross-validation within the training data, and a second validation with an independent, external-site data set. Models with the lowest error in predicting age on this validation set then served as candidates for the final model. Once the final model was determined, our hypotheses were then tested on the test set. Of importance, the test set was composed of untouched data that were not used to create, optimize or validate the model. Neither the model or the hypotheses were modified after the model was consider final and ready for hypothesis testing.

First, in order to reduce the number of inputs to the model and avoid circularity bias, we searched for the graph metrics that were the most reliably predictive of age.^4^ To do so, the training set data was entered in a support vector machine (SVM) and a regression tree ensemble model to estimate which graph metrics were the most important to predict chronological age (*i.e.*, highest weights). For the SVM model, the features (*i.e.*, the 26 metrics) were standardized by subtracting the mean and dividing by the standard deviation of the training group. SVM was implemented with the *fitrlinear* function using a linear kernel, and Bayesian-optimized ridge regularization. For ensemble methods, feature standardization is not recommended, and thus the unstandardized 26 metrics were used as input. The *fitrensemble* function was used with Bayesian optimization of hyperparameters including the method (‘Bag’ or ‘LSBoost’), number of learning cycles, and the learning rate. In both models, chronological age was the response vector, and parameter optimization was determined with the minimum 5-fold cross validation loss. Graph metrics were then ranked separately by order of SVM weights and ensemble model importance (i.e., highest load corresponding to the most important). “Importance” in the ensemble model was determined using the predictorImportance function, which is equal to the sum of changes in mean squared error due to splits on every predictor, divided by the number of branch nodes. The average rank from both models was then used to determine the overall importance of each metric.

In a second step, we aimed at creating an accurate model requiring the fewest number of features possible. We used training data to generate a neural net model and assessed its accuracy using the validation set. More specifically, the neural network was optimized by i) generating different models using the training set, each model varying in number of features used as input and network complexity, and ii) applying each model to the validation set (independent data set/site) to evaluate which one provided the better generalisability (*i.e.*, avoid overfitting and give the better prediction on an independent set). Graph metrics in both training and validation sets were standardized by subtracting the training group mean and dividing by the training group standard deviation. Network models had 5 to 25 input features in increments of 5, entered according to their importance, as determined previously (see above). A null model was also tested by applying the same feature increment procedure but entering the graph metrics in a random order. Architecture of the network was also tested with various number of hidden layers (1 or 2) and number of units in the hidden layers (2, 5, 7, or 10). Age was modeled on the training data using the *fitnet* function with Bayesian regularization backpropagation. Model accuracy was ultimately determined by the root-mean squared error (rmse) between actual and predicted age on the validation data; lower rmse reflecting higher accuracy. Because neural network units are initialized with random values, the rmse changed slightly each time model error was measured. Thus, the best model was determined by the lowest rmse, averaged over three iterations. Once the most accurate validated model was determined, it was applied on untouched data (test set).

### Additional measures in DIAN and PREVENT-AD samples (test set)

#### Genetics

DIAN genotyping was performed by the DIAN Genetics Core at Washington University and methods are fully described elsewhere.^19,80^ Briefly, the presence or absence of an ADAD mutation was determined using PCR-based amplification of the appropriate exon followed by Sanger sequencing. *APOE* genotype was determined using an ABI predesigned real-time Taqman assay.

*APOE* genotype in the PREVENT-AD was determined using the PyroMark Q96 pyrosequencer (Qiagen, Toronto, ON, Canada) and the following primers: rs429358_amplification_forward 5’-ACGGCTGTCCAAGGAGCT G-3’, rs429358_amplification_reverse_biotinylated 5’-CACCTCGCCGCGGTACTG-3’, rs429358_sequencing 5’-CGGACATGGAGGACG-3’, rs7412_amplification_forward 5’-CTCCGCGATGCCGATGAC-3’, rs7412_amplification_reverse_biotinylated 5’-CCCCGGCCTGGTACACTG-3’ and rs7412_sequencing 5’-CGATGACCTGCAGAAG-3’.

#### PET acquisition and processing

In DIAN, 28 mutation non-carriers and 117 mutation carriers from the test set had an Aβ-PET scans available at baseline. Aβ-PET scans were acquired in different centers, following ADNI protocol.^81^ Briefly, participants were injected intravenously with 8 mCi to 18 mCi of ^11^C-PIB. Part of the participants underwent a full dynamic acquisition of 70 minutes, starting at the time of injection. The remaining part of the sample underwent a 30-minute scan after a rest period of 40 minutes. A standard brain transmission scan (or computed tomography [CT] transmission scan for PET/CT scanners) was obtained for attenuation correction. Aβ-PET data was processed as previously described elsewhere.^82^ Standardized uptake value ratio (SUVR) were calculated using the cerebellar grey as a reference and a global measure of Aβ burden was calculated by averaging SUVRs from the prefrontal cortex, temporal lobe, gyrus rectus, and precuneus. A threshold of 1.31 was used to determine Aβ-positivity.^82,83^

In the PREVENT-AD cohort, Aβ-PET scans were performed at the MNI (Montréal, Canada) on a Siemens HRRT. Sixty-four individuals from the test set underwent this examination, at a mean distance of 10.30±5.63 months from their closest MRI session and 43.10±17.95 months after their baseline session. A 30-minute acquisition scan started 40 minutes after intravenous injection of approximately ~5.4mCi of ^18^F-NAV4694. Transmission scans were acquired for attenuation correction. Data was processed using a classic pipeline (see ^84^ and https://github.com/villeneuvelab/vlpp for details). A global index of neocortical Aβ burden was derived by extracting, in native space, the mean standardized uptake value ratio (SUVR) of the frontal, temporal, parietal and posterior cingulate cortex of the Desikan-Killiany atlas,^85^ using the cerebellum grey matter as reference region. A threshold for positivity was determined using Gaussian Mixture modelling^84^ and scans with global neocortical Aβ burden ≥ 1.39 were considered positive.

### Statistical analyses on the predicted age difference (test set)

To analyze the specificities of brain aging in the context of preclinical AD, we calculated the predicted age difference for DIAN and PREVENT-AD participants in the test set, as previously detailed,^3^ by subtracting the actual chronological age from the predicted brain age (output from the model). We were particularly interested in the influence of the genes involved in AD, which are either responsible of ADAD or increase the risk of sAD. To do so we compared, in the test set, the predicted age difference (i.e. PAD) between mutation non-carriers and mutation carriers from DIAN, and *APOE*4 carriers *vs* non-carriers in the PREVENT-AD. We were also interested to further understand the influence of Aβ accumulation on functional brain aging in asymptomatic individuals. To do so we assessed the effect of Aβ deposition, measured by PET, on the predicted age difference in both the DIAN and PREVENT-AD cohorts, both by comparing Aβ+ and Aβ-individuals (dichotomous variable), and a by assessing the correlation between predicted brain age difference and Aβ load (continuous variable). When applicable, post-hoc comparisons were done using Fisher’s LSD.

Analyses were conducted using Statistical Package for the Social Sciences (SPSS), and statistical significance was set at p<.05.

## Acknowledgments

The authors would like to thank the members of the Villeneuve Lab, J. Tremblay-Mercier, A. Labonté, D. Dea, C. Madjar and all the PREVENT-AD center for participants’ recruitment, data acquisition and data management (A complete listing of PREVENT-AD Research Group can be found in the PREVENT-AD database: https://preventad.loris.ca/acknowledgements/acknowledgements.php?date=[2019-07-30].); the members of the Brain Imaging Center of the Douglas Mental Health Institute for MRI acquisitions; the member of the Cyclotron and PET Units of the Montreal Neurological Institute for PET tracer production and acquisitions; K. Paumier, R. Hornbeck, P. Wang and S. Flores for their help in DIAN data preparation as well as all the centers involved in DIAN data acquisitions. This manuscript has been reviewed by DIAN Study investigators for scientific content and consistency of data interpretation with previous DIAN Study publications. We acknowledge the altruism of the participants and their families and contributions of the DIAN research and support staff at each of the participating sites for their contributions to this study.

Finally, we would like to thank all the participants and their family for their invaluable help.

