## Appendix for "Functional brain age prediction suggests accelerated aging in preclinical familial Alzheimer’s disease, irrespective of fibrillar amyloid-beta pathology"

Appendix – DIAN Study Group

| Last Name | First | Institution | Affiliation | Core | Role | Email address |
|---|---|---|---|---|---|---|
| Allegri | Ricardo | FLENI | FLENI Institute of Neurological Research (Fundacion para la Lucha contra las Enfermedades Neurologicas de la Infancia) | N/A | PI | |
| Bateman | Randy | WU | Washington University in St. Louis School of Medicine | Admin | **Core Leader/PI/Chair** | |
| Bechara | Jacob | Sydney | Neuroscience Research Australia | N/A | Site Leader | |
| Benzinger | Tammie | WU | Washington University in St. Louis School of Medicine | Imaging | **Core Leader** | |
| Berman | Sarah | Pitt | University of Pittsburgh | N/A | PI | |
| Bodge | Courtney | Butler | Brown University-Butler Hospital | N/A | Site Coordinator | |
| Brandon | Susan | WU | Washington University in St. Louis School of Medicine | Admin / Clincal | Core Personnel | |
| Brooks | William (Bill) | Sydney | Neuroscience Research Australia | N/A | Site Coordinator | |
| Buck | Jill | IU | Indiana University | N/A | Site Coordinator | |
| Buckles | Virginia | WU | Washington University in St. Louis School of Medicine | Admin | Core Personnel | |
| Chea | Sochenda | Mayo | Mayo Clinic Jacksonville | N/A | Site Coordinator | |
| Chhatwal | Jasmeer | BWH | Brigham and Women's Hospital–Massachusetts General Hospital | N/A | PI | |
| Chrem | Patricio | FLENI | FLENI Institute of Neurological Research (Fundacion para la Lucha contra las Enfermedades Neurologicas de la Infancia) | N/A | Site Coordinator | |
| Chui | Helena | USC | University of Southern California | N/A | PI | |
| Cinco | Jake | UCL | University College London | N/A | Site Coordinator | |
| Cruchaga | Carlos | WU | Washington University in St. Louis School of Medicine | Genetics | **Core Co-Leader** | |
| Donahue | Tamara | WU | Washington University in St. Louis School of Medicine | N/A | Site Coordinator | |
| Douglas | Jane | UCL | University College London | N/A | Site Coordinator | |

| Last Name | First | Institution | Affiliation | Core | Role | Email address |
|---|---|---|---|---|---|---|
| Edigo | Noelia | FLENI | FLENI Institute of Neurological Research (Fundacion para la Lucha contra las Enfermedades Neurologicas de la Infancia) | N/A | Site Coordinator | |
| Erekin-Taner | Nilufer | Mayo | Mayo Clinic Jacksonville | N/A | *sub-I* | |
| Fagan | Anne | WU | Washington University in St. Louis School of Medicine | Biomarker | **Core Leader** | |
| Farlow | Marty | IU | Indiana University | N/A | PI | |
| Fitzpatrick | Colleen | BWH | Brigham and Women's Hospital-Massachusetts | N/A | Site Co-Coordinator | |
| Flynn | Gigi | WU | Washington University in St. Louis School of Medicine | Admin / Clinical | Core Personnel | |
| Fox | Nick | UCL | University College London | N/A | PI | |
| Franklin | Erin | WU | Washington University in St. Louis School of Medicine | Neuropath | Core Coordinator | |
| Fujii | Hisako | Japan | Osaka City University | N/A | Assistant/Coord | |
| Gant | Cortaiga | WU | Washington University in St. Louis School of Medicine | Admin / Clinical | Core Personnel | |
| Gardener | Samantha | Perth | Edith Cowan University, Perth | N/A | Site Coordinator | |
| Ghetti | Bernardino | IU | Indiana University | N/A | *sub-I* | |
| Goate | Alison | Icahn NY | Icahn School of Medicine at Mount Sinai | Genetics | **Core Co-Leader** | |
| Goldman | Jill | CU | Columbia University | N/A | Genetics Ethics | |
| Gordon | Brian | WU | Washington University in St. Louis School of Medicine | Imaging | Core Personnel | |
| Graff-Radford | Neill | Mayo | Mayo Clinic Jacksonville | N/A | PI | |
| Gray | Julia | WU | Washington University in St. Louis School of Medicine | Biomarker | Core Personnel | |
| Groves | Alexander | WU | Washington University in St. Louis School of Medicine | Biomarker | Core Coordinator | |
| Hassenstab | Jason | WU | Washington University in St. Louis School of Medicine | Clinical | Core Personnel | |
| Hoechst-Swisher | Laura | WU | Washington University in St. Louis School of Medicine | Admin / Clinical | Core Coordinator | |

| Last Name | First | Institution | Affiliation | Core | Role | Email address |
|---|---|---|---|---|---|---|
| Holtzman | David | WU | Washington University in St. Louis School of Medicine | N/A | Associate Director | |
| Hornbeck | Russ | WU | Washington University in St. Louis School of Medicine | Imaging | Core Coordinator | |
| Houeland DiBari | Siri | Munich | German Center for Neurodegenerative Diseases (DZNE) Munich | N/A | Site Coordinator | |
| Ikeuchi | Takeshi | Niigata | Niigata University | N/A | *Site Leader* | |
| Ikonomovic | Snezana | Pitt | University of Pittsburgh | N/A | Site Coordinator | |
| Jack | Clifford | Mayo | Mayo Clinic Jacksonville | MRI QC | Vendor MRI QC | |
| Jerome | Gina | WU | Washington University in St. Louis School of Medicine | Biomarker | Core Coordinator | |
| Jucker | Mathias | Tubingen | German Center for Neurodegenerative Diseases (DZNE) Tubingen | N/A | PI | |
| Karch | Celeste | WU | Washington University in St. Louis School of Medicine | Administrative | Core Personnel | |
| Kasuga | Kensaku | Niigata | Niigata University | N/A | Site Coordinator | |
| Kawarabayashi | Takeshi | Hirosaki | Hirosaki University | N/A | Clinician | |
| Klunk | William (Bill) | Pitt | University of Pittsburgh | N/A | sub-I | |
| Koeppe | Robert | U of Michigan | University of Michigan | PET QC | Vendor PET QC | |
| Kuder-Buletta | Elke | Tubingen | German Center for Neurodegenerative Diseases (DZNE) Tubingen | N/A | Site Coordinator | |
| Laske | Christoph | Tubingen | German Center for Neurodegenerative Diseases (DZNE) Tubingen | N/A | *sub-I* | |
| Lee | Jae-Hong | Korea | Asan Medical Center | N/A | PI | |
| Levin | Johannes | Munich | German Center for Neurodegenerative Diseases (DZNE) Munich | N/A | PI | |
| Martins | Ralph | Perth | Edith Cowan University | N/A | PI | |
| Mason | Neal Scott | UPMC | University of Pittsburgh Medical Center | PIB QC | Vendor PIB QC | |
| Masters | Colin | Melb | University of Melbourne | N/A | PI - former | |

| Last Name | First | Institution | Affiliation | Core | Role | Email address |
|---|---|---|---|---|---|---|
| Maue-Dreyfus | Denise | WU | Washington University in St. Louis School of Medicine | Clinical | Core Personnel | |
| McDade | Eric | WU | Washington University in St. Louis School of Medicine | Clinical | **Core Leader Assoc** | |
| Mori | Hiroshi | Japan | Osaka City University | N/A | PI | |
| Morris | John | WU | Washington University in St. Louis School of Medicine | Clinical | **Core Leader** | |
| Nagamatsu | Akem | Tokyo | Tokyo University | N/A | Site Coordinator | |
| Neimeyer | Katie | CU | Columbia University | N/A | Site Coordinator | |
| Noble | James | CU | Columbia University | N/A | PI | |
| Norton | Joanne | WU | Washington University in St. Louis School of Medicine | Genetics | Core Coordinator | |
| Perrin | Richard | WU | Washington University in St. Louis School of Medicine | Neuropath | **Core Leader** | |
| Raichle | Marc | WU | Washington University in St. Louis School of Medicine | Imaging | Core Personnel | |
| Renton | Alan | Icahn NY | Icahn School of Medicine at Mount Sinai | Genetics | Core Personnel | |
| Ringman | John | USC | University of Southern California | N/A | *sub-I* | |
| Roh | Jee Hoon | Korea | Asan Medical Center | N/A | *sub-I* | |
| Salloway | Stephen | Butler | Brown University-Butler Hospital | N/A | PI | |
| Schofield | Peter | Sydney | Neuroscience Research Australia | N/A | PI | |
| Shimada | Hiroyuki | Osaka | Osaka City University | N/A | *Site Leader* | |
| Sigurdson | Wendy | WU | Washington University in St. Louis School of Medicine | N/A | Site Coordinator | |
| Sohrabi | Hamid | Perth | Edith Cowan University | N/A | Site Coordinator | |
| Sparks | Paige | BWH | Brigham and Women's Hospital-Massachusetts | N/A | Site Coordinator | |
| Suzuki | Kazushi | Tokyo | Tokyo University | N/A | *Site Leader* | |
| Taddei | Kevin | Perth | Edith Cowan University | N/A | Site Coordinator | |
| Wang | Peter | WU | Washington University in St. Louis School of Medicine | Biostat | Core Coordinator | |
| Xiong | Chengjie | WU | Washington University in St. Louis School of Medicine | Biostat | **Core Leader** | |

| Last Name | First | Institution | Affiliation | Core | Role | Email address |
|---|---|---|---|---|---|---|
| Xu | Xiong | WU | Washington University in St. Louis School of Medicine | Biostat | Core Personnel | |
| Levey | Allan | Emory | Emory University School of Medicine | N/A | Project Leader | |
